# Quantitative reanalysis of classic pigeon homing experiments rejects a liver-based magnetic compass based on superparamagnetism

**DOI:** 10.64898/2026.09.07.749779

**Authors:** Joseph L. Kirschvink, Charles Walcott

## Abstract

Lisowski et al. [1] recently proposed that the pigeon’s magnetic compass resides in the liver and depends upon ferritin-containing macrophages operating through a superparamagnetic mechanism. Here we evaluate this hypothesis using behavioral, neurobiological, evolutionary, and biophysical evidence together with three original analyses. First, we present a quantitative reanalysis of the classic pigeon homing experiments of Walcott and Green [2], calculating for the first time the magnetic fringe-field distribution produced by the head-mounted coil apparatus. These calculations demonstrate that the liver would have experienced a maximum shift of about 1.4% of the ambient geomagnetic field, far too small to account for the observed orientation effects, thereby localizing the relevant magnetoreceptive structures to the region of the head and excluding the liver. Second, we quantify the magnetic alignment expected for ferritin cores under geomagnetic conditions and show that ferrihydrite nanoparticles are approximately five orders of magnitude too weak to resist thermal noise, while their superparamagnetic moments fluctuate on timescales roughly eleven orders of magnitude faster than those relevant to navigation. Third, we analyze the neurobiological requirements of magnetic navigation and show that mobile macrophages are unlikely to provide stable directional information because compass signals must be continuously cross-calibrated against vestibular and visual inputs within a body-centered reference frame. Comparative evidence from fish, birds, and turtles further associates magnetically responsive structures with craniofacial tissues and trigeminal sensory pathways. We suggest the disruption of the magnetic response is due to induced anemia from clodronate treatment that triggers dissolution and absorption of iron in the magnetite-containing cells involved in magnetoreception. Together these findings indicate that a liver-based ferritin compass is inconsistent with behavioral, neurobiological, and physical evidence and instead support cephalized magnetite-based receptor systems as the basis of avian magnetoreception.

## Introduction

More than fifty years ago, Walcott and Green [2] demonstrated that magnetic fields applied locally to the heads of homing pigeons predictably altered orientation behavior under overcast conditions. Battery-powered coils mounted around the head generated magnetic fields comparable to the ambient geomagnetic field. When the magnetic field experienced by the head was inverted upwards, pigeons departed in directions approximately opposite to the homeward bearing, a finding consistent with the known inclination-dependent magnetic compass in birds.

However, a recent report by Lisowski et al. [1] in the Journal *Science* proposed that the pigeon’s magnetic compass resides in the liver and is mediated by ferritin-containing macrophages whose superparamagnetic properties provide directional information. If correct, this interpretation would require major revision of prevailing models of vertebrate magnetoreception, which have generally focused on cephalized receptor systems with magnetite-containing structures associated with the trigeminal nerve, and/or on possible quantum effects on pigments in the retina.

Several independent lines of evidence are relevant to evaluating this macrophage hypothesis. As noted above, classic behavioral studies demonstrated that magnetic manipulations applied to the head alter homing orientation in pigeons, but Walcott and Green [2] did not evaluate the effect of those coils at the position of the liver. Neurobiological investigations across vertebrates have also localized putative magnetoreceptive structures to craniofacial tissues innervated by trigeminal pathways [3-6]. Finally, physical analyses of candidate receptor materials indicate that magnetite-based mechanisms have properties consistent with geomagnetic sensing, whereas superparamagnetic mechanisms of any sort face substantial thermodynamic limitations [7].

Importantly, this study is not solely a review of prior work. We present three original analyses that directly test central predictions of the liver-compass hypothesis. First, we reconstruct the magnetic fringe-field distribution generated by the head-mounted coil apparatus of Walcott and Green [2], providing a quantitative test of whether alteration of the field at the position of the liver could have mediated the observed behavioral effects. Second, we evaluate ferritin alignment, thermal-noise limits, and magnetic relaxation under geomagnetic conditions to assess the physical feasibility of the proposed superparamagnetic mechanism. Third, we analyze the neurobiological requirements of multisensory compass calibration and ask whether mobile macrophages could provide stable directional information to the brain. Together, these analyses test the anatomical localization, physical plausibility, and neural functionality of the proposed liver-based compass. All tests fail, and we provide a simpler hypothesis for the Lisowski et al. [1] results that support a magnetite-based magnetoreception mechanism in the head.

## Methods

As the displacement homing experiments employed by Lisowski et al. [1] involve the same comparison of homing ability on sunny vs. cloudy days that was employed by Walcott and Green [2], it is a good assumption that both experiments activated the same magnetoreceptive sensory organs. Hence, the hypothesis that these magnetoreceptors might be in the liver can be tested by calculating the fringe field of the paired Walcott and Green [2] coils that were wrapped on the bird’s head as shown here in Fig. 1. These two coils were not a classical Helmholtz pair, so it is necessary to calculate the fringe fields produced by each coil at the position of the pigeon’s liver, and do the vector sum of the two contributions. We consider two cases, one when the bird has its head aligned so the liver is on the axis parallel to the plane of the coils, and the other when the head is tilted 90° so the liver is located perpendicular to the coil axis (red dotted outlines as shown in Fig 1). A bird’s head is highly mobile, so both orientations need consideration and these represent the two end-member cases of all possible head orientations. The liver is ∼100 mm from the head center in both orientations (CW, personal observations); however, this distance is not critical as the far-field effect drops off as the inverse cube of the distance. Data from CW’s notebooks and from Walcott and Green [2] indicate that the coils were approximately co-planer, wired in series, each wrapped with 200 turns of fine-gauge magnet wire, had respective diameters of 23 and 35 mm, and were separated by 31 mm (see Fig. 1). They measured the field produced in the center of the coil pair (which would have been located inside the pigeon’s head when it was mounted) at ∼ 60 µT (0.6 G) with a Hall probe. With this geometry, and as outlined on Table 1 and the Supplemental Information, we can back-calculate the effective current flowing through each loop, and thereby the magnitude of the field at all points in space surrounding the coil pair, including at the position of the liver.

**Table 1.**
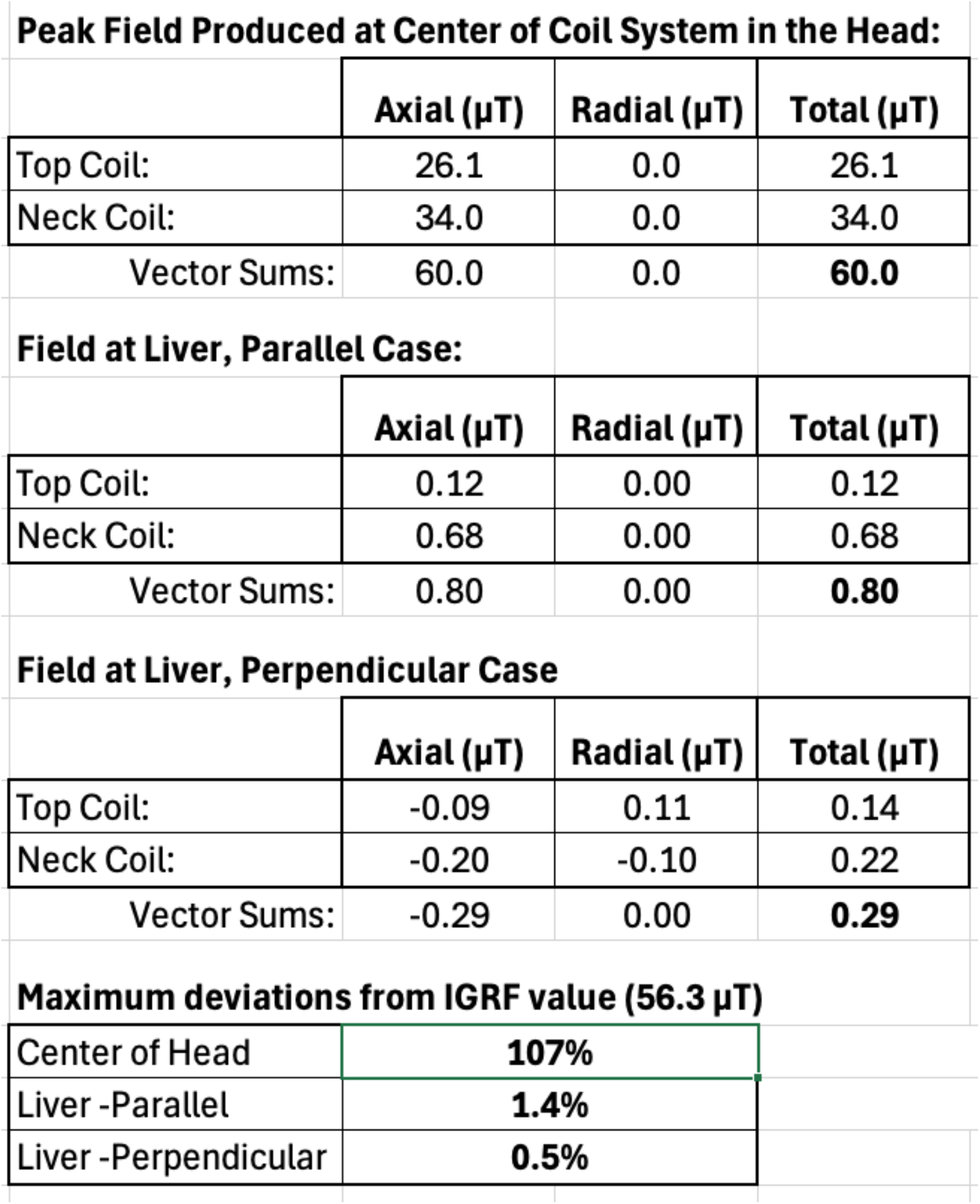
Quantitative reconstruction of magnetic fringe-field exposure at the liver during the Walcott and Green homing experiments. Calculated magnetic field intensities at the liver for the on-axis (parallel) and off-axis (perpendicular) coil orientations shown in Figure 1. Calculations assume a distance of approximately 10 cm from the coil center to the nearest edge of the liver. Axial and off-axis field components were calculated using complete elliptic-integrals of the first and second kinds for circular current loops [8, 9]. Geomagnetic reference values for Stony Brook, New York, were obtained from International Geomagnetic Reference Field (IGRF) models. This analysis demonstrates that the liver would have experienced only a small fraction of the ambient geomagnetic field despite strong behavioral effects produced by the head-mounted coils, providing a direct quantitative test of the liver-localization hypothesis. To our knowledge, these are the first calculations of the magnetic fringe fields experienced by the liver during the Walcott and Green experiments and provide a direct quantitative evaluation of the liver-compass hypothesis.

**Peak Field Produced at Center of Coil System in the Head:**
|  | **Axial ($\mu\text{T}$)** | **Radial ($\mu\text{T}$)** | **Total ($\mu\text{T}$)** |
| --- | --- | --- | --- |
| Top Coil: | 26.1 | 0.0 | 26.1 |
| Neck Coil: | 34.0 | 0.0 | 34.0 |
| Vector Sums: | 60.0 | 0.0 | **60.0** |

**Field at Liver, Parallel Case:**
|  | **Axial ($\mu\text{T}$)** | **Radial ($\mu\text{T}$)** | **Total ($\mu\text{T}$)** |
| --- | --- | --- | --- |
| Top Coil: | 0.12 | 0.00 | 0.12 |
| Neck Coil: | 0.68 | 0.00 | 0.68 |
| Vector Sums: | 0.80 | 0.00 | **0.80** |

**Field at Liver, Perpendicular Case**
|  | **Axial ($\mu\text{T}$)** | **Radial ($\mu\text{T}$)** | **Total ($\mu\text{T}$)** |
| --- | --- | --- | --- |
| Top Coil: | -0.09 | 0.11 | 0.14 |
| Neck Coil: | -0.20 | -0.10 | 0.22 |
| Vector Sums: | -0.29 | 0.00 | **0.29** |

**Maximum deviations from IGRF value (56.3 $\mu\text{T}$ )**
|  |  |
| --- | --- |
| Center of Head | **107%** |
| Liver -Parallel | **1.4%** |
| Liver -Perpendicular | **0.5%** |

**Figure 1.**
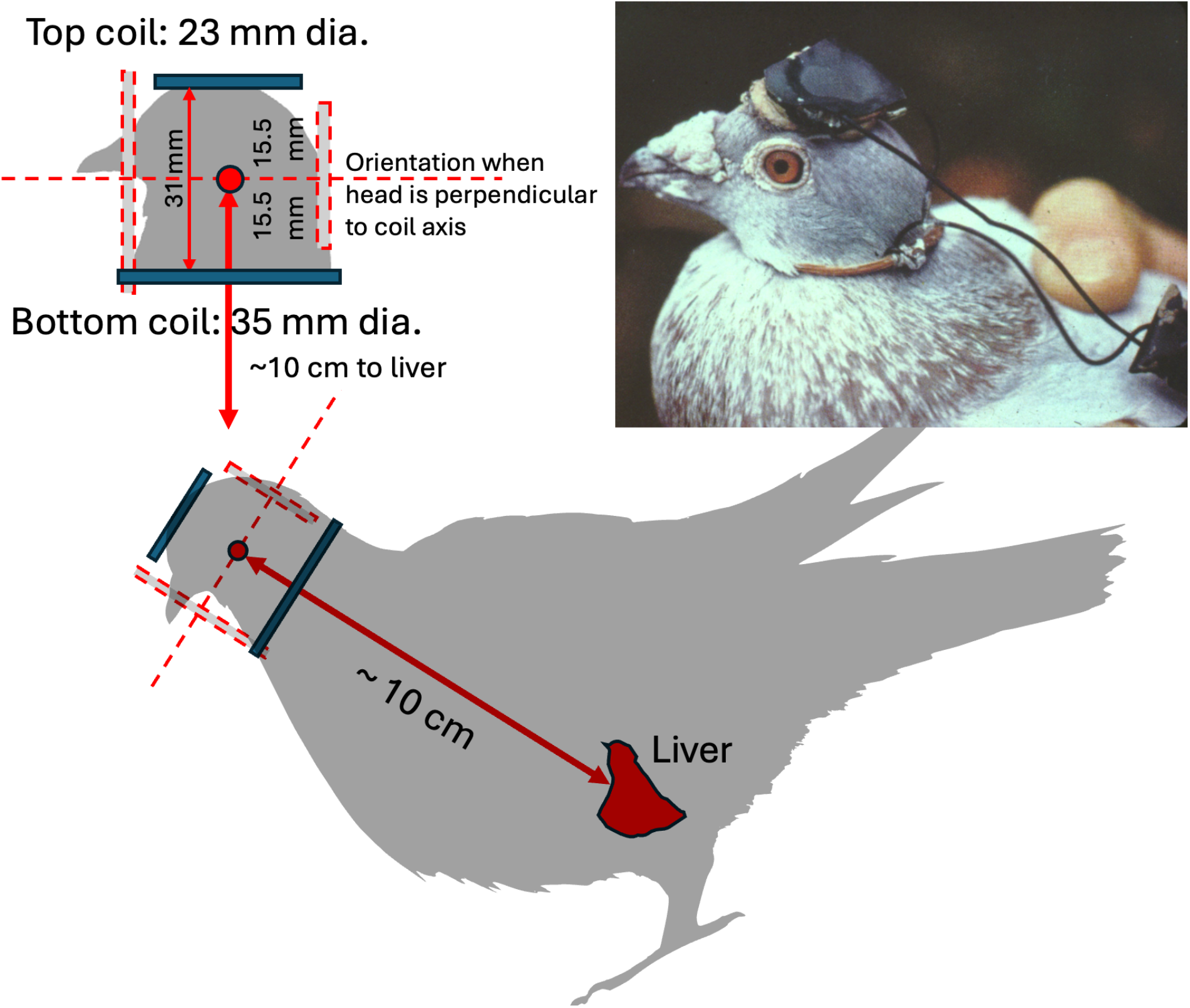
Quantitative reanalysis of the head-mounted coil experiment of Walcott and Green. Schematic diagram of the head-mounted coils used by Walcott and Green [2], superimposed to scale on a pigeon. Two orientations are shown because of head mobility: one with the liver approximately aligned with the coil axis (solid lines) and one approximately perpendicular to it (dashed lines). The geometry illustrated here was used for the magnetic field calculations presented in Table 1 and Supporting Online Information. These calculations constitute a new quantitative analysis of the original experimental apparatus and were performed to test whether magnetic stimulation of the liver could plausibly account for the behavioral effects reported by Walcott and Green. Inset: photograph of a pigeon carrying the coil apparatus, taken by CW in 1973. Pigeon silhouette adapted from “Pigeon silhouette 4874.svg” by Dori and Nevit Dilmen, Wikimedia Commons, CC BY-SA 3.0. Liver position added and modified by the authors.

The calculation of the field produced along the axis of a circular coil has analytical solutions [8], as noted in the Supplemental Information. Using this, and the configuration above, we find that a net current of 2.255 Amperes will produce a 60 µT field in the coil center, or a net current of 11 mA through the 200 turns of wire in each coil of the pair. This will suffice for the on-axis case where the head is positioned such that the liver is in-line with the coils, but the bird’s head can move in nearly all directions. We therefore need to use the more complete off-axis solutions that involve the use of the Biot-Savart law, with the extreme case when the liver is aligned perpendicular to the coil axes (red dotted lines in Fig. 1). This requires calculation of complete elliptical integrals of the first and second kind, as is standard practice [8, 9]. Results for each coil in the two orientations are shown in Table 1, with the calculations explained in the supplemental Excel sheet.

Calculations on the possibility that the superparamagnetic properties of aggregated ferritin cores might provide the biophysical basis of a magnetoreceptor cell were done using the well-known properties of ferritin [10], coupled with standard statistical mechanics and use of the Langevin function for the alignment of magnetic moments against thermal agitation, as was done for the magnetotactic bacteria over 45 years ago [11].

## Results

### Quantitative Reanalysis of the Classic Homing Experiments of Walcott & Green [2]

The liver-compass hypothesis proposed by Lisowski et al. [1] makes a straightforward prediction: if liver-associated receptors mediated the behavioral effects observed by Walcott and Green [2], then the magnetic fields generated by the head-mounted coil apparatus must have substantially affected the liver.

To test this prediction, we performed new calculations of the magnetic field distribution generated by the experimental coils [see Methods]. These calculations were not presented in the original Walcott and Green [2] study and constitute an original component of the present work.

Using the geometry of the original apparatus (Fig. 1), we calculated the magnetic fringe fields extending from the coils to the liver. Assuming a distance of approximately 10 cm from the coil center to the nearest edge of the liver, the maximum field strength at the liver when the head is aligned with the liver on the axis of the coils would have been only about 1.4% of the ambient geomagnetic field (Table 1; Supporting Online Information). The perpendicular alignment yields a maximum deviation of only 0.5%.

Our analysis demonstrates that the liver was exposed to only a weak residual fringe field despite profound behavioral effects produced by the coils. The field at the compass receptors needs to have the vertical component inverted in order to elicit the ∼180° change in homing direction observed by Walcott and Green [2]. Consequently, their results cannot plausibly be attributed to stimulation of receptors in the liver. Instead, these experiments provide direct quantitative evidence that the relevant magnetoreceptive structures are in the vicinity of the head.

### Quantitative Evaluation of Ferritin-Based Magnetoreception

We next examined whether ferritin mineralization could plausibly function as the physical basis of a geomagnetic compass, as claimed by Lisowski et al. [1]. Using published magnetic properties of ferrihydrite and standard thermal-noise calculations [10], we estimated the expected magnetic alignment of ferritin particles under geomagnetic field strengths.

Ferritin cores consist primarily of ferrihydrite, a poorly crystalline iron oxyhydroxide with relatively weak magnetic properties. Although ferrihydrite nanoparticles can exhibit minor ferrimagnetic contributions arising from structural defects and uncompensated electron spins at the crystal surface, their saturation magnetization remains far below that of magnetite or maghemite [10].

An 8-nm ferrihydrite particle possesses a magnetic moment, µ, of only about 1 × 10^−21^ A·m^2^. Under geomagnetic conditions this corresponds to a magnetic-to-thermal energy ratio (µB/kT) of approximately 1.2 × 10^−5^ (where B is the Earth’s field strength, k is the Boltzmann constant, and T the absolute temperature), yielding a predicted magnetic alignment (given by the Langevin function) of only 0.0004%.

Thus, ferritin particles are approximately five orders of magnitude too weak to maintain biologically meaningful alignment against thermal fluctuations. Aggregates of ferritin cores do not do much better.

A second limitation involves magnetic relaxation. Because ferritin particles are superparamagnetic, their magnetic moments are not rigidly locked to crystal orientation and therefore cannot generate substantial mechanical torques. Furthermore, ferritin cores undergo Néel-Brown magnetic fluctuations on nanosecond timescales [12]. These fluctuations occur approximately eleven orders of magnitude faster than the behavioral timescales relevant to navigation proposed by Lisowski et al [1].

The proposed ferritin-based mechanism therefore lacks both the strength and temporal stability required for directional sensing.

## Discussion

### Original Contributions of the Present Study

Although motivated by the conclusions of Lisowski et al. [1], this manuscript is not simply a review or commentary. Rather, it provides three independent analyses that directly test critical assumptions of the liver-compass hypothesis.

First, we calculated the magnetic fringe fields experienced by the liver during the classic Walcott and Green homing experiments [2]. These calculations show that the liver would have experienced only a small fraction of the ambient geomagnetic field despite profound behavioral effects produced by magnetic stimulation of the head.

Second, we quantified the expected magnetic alignment and relaxation behavior of ferritin particles under geomagnetic conditions. These calculations demonstrate that ferritin-based superparamagnetism is both too weak and temporally too unstable to serve as the basis of a biological compass.

Third, we analyze below the neurobiological requirements of multisensory calibration and show that mobile macrophages are unlikely to provide the stable body-centered reference signals required for directional navigation. A magnetic compass must remain calibrated to vestibular and visual systems over biologically relevant timescales, a requirement difficult to reconcile with a receptor system built from migratory immune cells.

These anatomical, biophysical, and neurobiological analyses are independent of one another, yet all converge upon the same conclusion that the present study identifies fundamental difficulties with both the proposed anatomical location and the proposed physical mechanism of the liver-based compass model.

### Other Comparative Neurobiological Evidence for Craniofacial Magnetoreceptors

Evidence from other vertebrates similarly localizes magnetoreception to the craniofacial region.

Walker et al. [3] and Diebel et al. [4] traced magnetically responsive pathways associated with the rostral ophthalmic branch of the trigeminal nerve in trout. These structures were associated with single-domain ferromagnetic material and occurred in tissues previously shown to contain biogenic magnetite by SQUID magnetometry [13], and extraction followed by transmission electron microscopy revealed chains of single-domain magnetite crystals similar in form and organization to those from magnetotactic bacteria [14].

Magnetically responsive neurons have also been identified in the ophthalmic branch of the trigeminal nerve in birds [5], including pigeons [6]. Importantly, these pathways innervate the head rather than visceral organs such as the liver.

More recently, studies of hatchling sea turtles have similarly constrained magnetite-based receptor structures to the head region [15].

Collectively, these observations indicate that vertebrate magnetic sensory systems are consistently associated with craniofacial tissues and trigeminal pathways rather than abdominal organs.

### Multisensory Calibration Requires Stable Magnetoreceptors

A magnetic compass is not merely a magnetic detector. To generate biologically useful directional information, magnetic signals must be integrated continuously with gravitational, vestibular, and visual inputs from the animal and processed in the brain.

Because magnetic and gravitational fields are physically independent, the nervous system must learn how signals from magnetoreceptors covary with head and body movements. A magnetoreceptor cell sending a modulated stream of action potentials to the brain cannot also include vestibular information in that data stream. Information about magnetic declination and inclination can only be obtained by having the brain cross-calibrate the magnetic data stream at the level of individual cells with the vestibular and gravitational systems. Such multisensory calibration is the only path for extracting inclination and compass-direction information from environmental magnetic fields [16].

This requirement imposes a fundamental constraint on receptor design. Magnetoreceptors must remain sufficiently stable relative to the body that their outputs can be interpreted within a consistent spatial reference frame. Long-term calibration becomes difficult if receptor locations and orientations continuously change.

Macrophages appear poorly suited to satisfy this requirement. These cells migrate, alter their morphology, and remodel their local interactions with surrounding tissue. Even if a macrophage contained magnetic material and occurred near sensory fibers, its movement would continuously alter the spatial relationship among magnetic particles, neighboring cells, and afferent neural pathways. Consequently, the resulting signal would depend partly on the changing position and orientation of the macrophage itself.

To our knowledge, this issue has not been explicitly addressed in models of magnetoreception. Yet a directional compass requires stable neural reference signals over timescales sufficient for learning, calibration, maintaining synaptic contacts, and navigation. The mobility and plasticity of macrophages appear fundamentally incompatible with these requirements.

This need for multisensory integration further suggests that evolution should favor anatomically stable and cephalized magnetic sensory systems, consistent with the organization observed for other major sensory modalities in bilaterian animals [17].

### Alternative Interpretation of the Clodronate Experiments

Macrophages play a central role in systemic iron metabolism [18]. Clodronate liposome treatment can disrupt macrophage-mediated iron recycling and can induce anemia and other alterations in iron homeostasis [19].

A simpler interpretation of the loss of homing ability reported following clodronate treatment is that disruption of iron metabolism adversely affects magnetite-based receptors located elsewhere in the body, including candidate receptor structures within the head. Iron is essential for numerous fundamental biochemical reactions, so it is not unreasonable that a global anemic stress might activate siderophore systems able to mine the iron from magnetite-based sensory cells in the region of the head.

Under this interpretation, the observed behavioral effects would support rather than contradict magnetite-based models of magnetoreception [7, 20].

## Conclusions

1. New magnetic-field calculations presented here demonstrate that the head coils used in the classic Walcott and Green homing experiments [2] would have produced only a small alteration of the ambient geomagnetic field at the position of the liver.
2. This quantitative reanalysis provides a direct test of the liver-localization hypothesis and indicates that the behavioral effects originated from structures located in the head.
3. Comparative evidence from fish, birds, and turtles consistently localizes candidate magnetoreceptive structures to craniofacial regions and trigeminal pathways.
4. Directional magnetic sensing requires stable calibration of the magnetic with vestibular and visual systems. The mobility and continual remodeling of macrophages make them poorly suited to provide the persistent body-centered reference signals expected of a biological compass receptor.
5. Ferritin-based superparamagnetism is approximately five orders of magnitude too weak and eleven orders of magnitude too dynamically unstable to function as a biologically useful magnetic compass.
6. The effects of clodronate treatment may reflect disruption of iron metabolism and mining of iron from magnetite-based receptor cells rather than elimination of macrophage-based magnetoreceptors.
7. The present study contributes three original analyses, anatomical, biophysical, and neurobiological, that independently reject a liver-based ferritin compass and strengthen the case for cephalized magnetite-based magnetoreception in birds.

## Supporting information

Supplemental Excel Sheet

## Supplemental Information

An Excel sheet, *Pigeon_Coil_and_Thermal_Noise_Calculations_Matching_Walcott&Green*.*xlsx*, with 3 tabs that summarize the data in Table 1, the magnetic field calculations, and the magnetic alignment of the ferritin cores with respect to thermal noise, respectively, has been uploaded as supplemental information.

### Competing interests

The authors declare no competing financial or non-financial interests.

Submission Notice: An earlier version of this ms was originally submitted as a formal ***Letter*** to **Science** on 21 July 2026 (manuscript aek7761). We considered this the appropriate venue because the Lisowski et al. paper was published as a full Research Article, accompanied by a News item, and featured on the cover of the 28 May issue.

In our view, the paper contains fundamental flaws and advances conclusions that set the science of magnetoreception back by more than 50 years. Accordingly, we believed that a formal, peer-reviewed response was warranted.

However, the editors of Science declined to send our submission for review and instead suggested that we post a 300-word response as an eLetter. Because eLetters are neither peer-reviewed nor citable publications, we do not regard that format as an adequate mechanism for addressing substantive scientific concerns. We will post a brief note there linking to this bioRxiv manuscript but are seeking publication of this critique in a journal with higher standards of scientific review and discourse.

## Notes

### Competing Interest Statement

The authors have declared no competing interest.

